# Impedance scaling for gold and platinum microelectrodes

**DOI:** 10.1101/2021.03.09.434678

**Authors:** Bo Fan, Bernhard Wolfrum, Jacob T. Robinson

**Affiliations:** Department of Electrical and Computer Engineering, Rice University, Houston, Texas 77005, United States; Department of Electrical and Computer Engineering, Technical University of Munich, Garching, 85748, Germany; Department of Bioengineering, Rice University, Houston, Texas 77005, United States; Department of Neuroscience, Baylor College of Medicine, Houston, Texas 77030, United States

## Abstract

**Objective:** Electrical measurement of the activity of individual neurons is a primary goal for many invasive neural electrodes. Making these “single unit” measurements requires that we fabricate electrodes small enough so that only a few neurons contribute to the signal, but not so small that the impedance of the electrode creates overwhelming noise or signal attenuation. Thus, neuroelectrode design often must strike a balance between electrode size and electrode impedance, where the impedance is often assumed to scale linearly with electrode area.

**Approach and main results:** Here we study how impedance scales with neural electrode area and find that the 1 kHz impedance of Pt electrodes (but not Au electrodes) transitions from scaling with area (r^-2^) to scaling with perimeter (r^-1^) when the electrode radius falls below 10 microns. This effect can be explained by the transition from planar to spherical diffusion behavior previously reported for electrochemical microelectrodes. *Significance:* These results provide important intuition for designing small, single unit recording electrodes. Specifically, for materials where the impedance is dominated by a pseudo-capacitance that is associated with a diffusion limited process, the total impedance will scale with perimeter rather than area when the electrode size becomes comparable with the diffusion layer thickness. For Pt electrodes this transition occurs around 10 um radius electrodes. At even lower frequencies (1 Hz) impedance approaches a constant. This transition to r^-1^ scaling implies that electrodes with a pseudo-capacitance can be made smaller than one might expect before thermal noise or voltage division limits the ability to acquire high-quality single-unit recordings.

## Introduction

Neurons, as the fundamental functional units of the nervous network, communicate using electrophysiological signals^1^. This makes neural electrodes the most commonly used tools for numerous discoveries in neuroscience^2-5^ and clinical therapies^6,7^. Penetrating microelectrodes can be implanted deep in the brain for extracellular recording from nearby neurons. Extracellular recordings are a mixture of local-field potential (LFP) and single-unit recording, which can be separated by filtering. The filtering bandwidth is 300-5000 Hz for single unit recording and < 300 Hz for LFP signals. Stable, chronic and high-quality single unit recording is essential to understand the neural networks on cellular levels^8,9^, and it provides the most precise control in brain-machine interface applications^6,7^.

Small electrodes are desirable for recording the activity of individual neurons (or “single units”) because their high spatial resolution allows them to record higher amplitude signals compared to large electrodes^10^. In addition, for penetrating electrodes, smaller footprints reduce the damage to neural tissue upon implantation. On the other hand, smaller electrodes also have a higher impedance, which can reduce the quality of neural recordings. This high impedance can decrease the signal-to-noise ratio (SNR) in two ways. First, it leads to higher thermal noise, which increases for voltage recordings as impedance increases^11,12^. Second, it creates a voltage divider, which consists of the electrode and the recording system in series^11-13^. As a result, higher impedance above 2-5 MΩ^11,13^ can cause greater signal attenuation, depending on the design of the neural probes and amplifier system.

The important trade-offs in performance as a function of electrode size, makes it important to understand how the impedance scales with the electrode area, particularly for ultra-small electrodes. For planar electrodes it is typically assumed that impedance is inversely proportional to the electrode area^12,14^ and thus electrode materials are often characterized by a specific impedance (impedance times area) that is assumed to be a constant for that material. For volumetric conductors, like poly(3,4-ethylenedioxythiophene)(PEDOT) the impedance scales inversely with the volume of the electrode coating^15^. For planar metallic electrodes, a systematic study of impedance scaling in Pt, gold and PEDOT coated electrodes with radii ranging from 10 - 1000 um concluded that the porous coatings contribute to lower charge transfer resistance and better capacitive coupling of the electrode^16^.

Recently we noted that impedance scales differently in flat and porous Pt of different areas and that the perimeter of electrodes might also affect the impedance^12^. Here, we systematically study the impedance of Pt and Au electrodes as a function of size from radius ranging from 3 - 125 µm to see how impedance scales with electrode dimensions and what mechanism may explain the transition between area dependence and perimeter dependence.

## Results

To evaluate the scaling of electrode impedances we first fabricated raised Pt microelectrodes with radius ranging from 3 to 125 µm using standard microfabrication techniques^12^ (see Methods). Briefly we fabricated 16-channel flexible microelectrodes using SU8 as an insulation layer. All 16 channels, though exposed electrodes are of different sizes, share one large via on the top SU8 for the raised structure (figure 1a).

**Figure 1.**
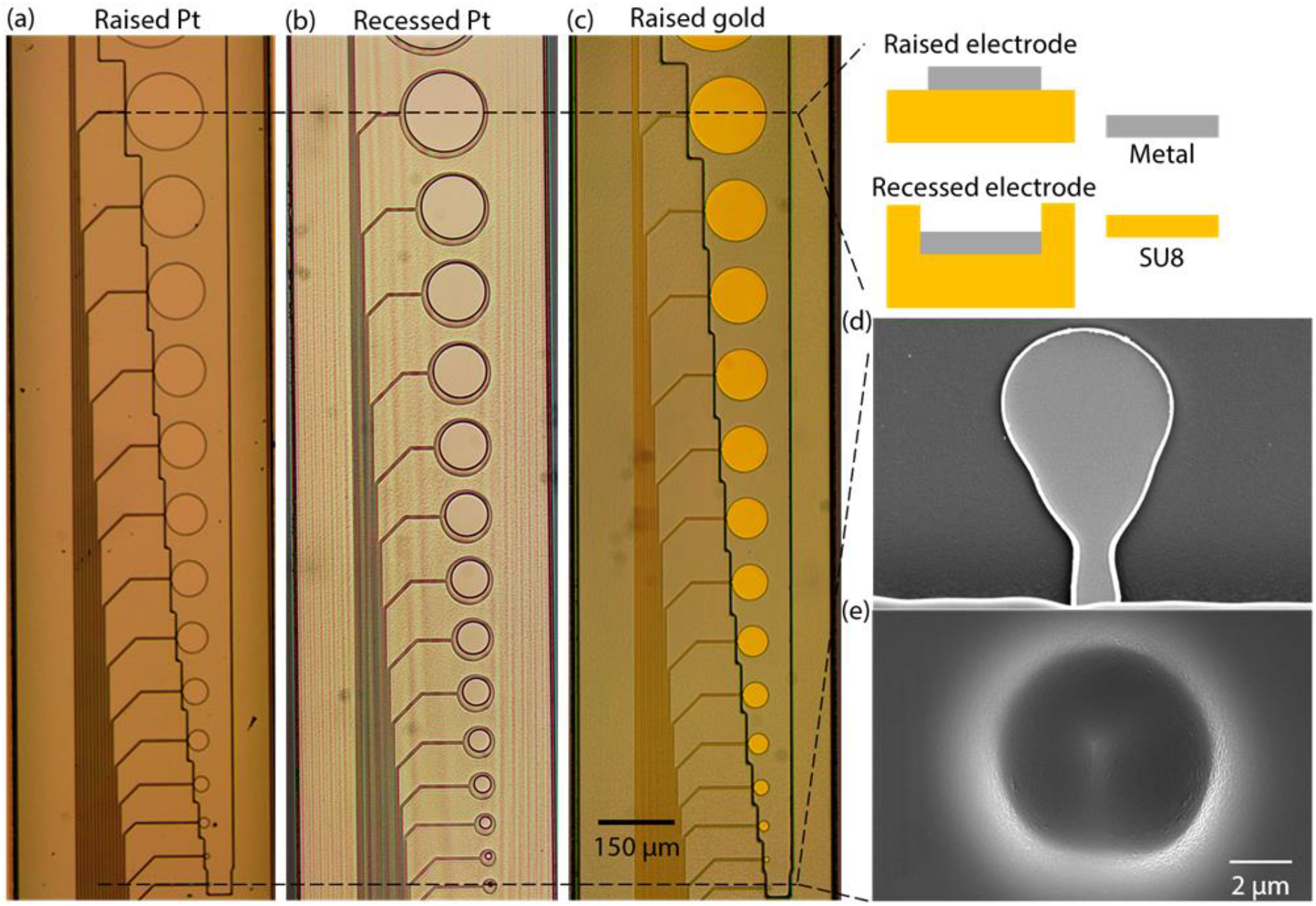
Microscopic image and SEM image of raised Pt, recessed Pt and raised Au electrodes. (a)-(c) 5X microscopic image of raised Pt, recessed Pt and raised Au electrodes, respectively. Inlet: Schematic of raised electrodes (top) and recessed electrodes (bottom). (d) SEM image of a raised Pt electrode with a radius of 3 µm. (e) SEM image of a recessed Pt electrode with a radius of 3 µm.

We then measured the 1 kHz impedance and electrochemical impedance spectra (EIS) for each differently sized electrode (SI figure 1a,1d). For these measurements we used a Gamry 600+ with a three-electrode setup in 1X phosphate-buffered saline (PBS) solution with a pH of 7.4. The input AC signal is 10 mV in root-mean-square to ensure a pseudo-linear relationship between current and voltage for EIS measurements using Ohm’s Law. We used Ag/AgCl as the reference electrode (0.199 V vs. standard hydrogen electrode) and a bare Pt wire 0.010’’ in diameter as the counter electrode (For more details see Methods).

We plotted these impedance data at 1 kHz versus radius on a log scale (figure 2a). For larger electrodes (radius r >= 10 µm), the impedance has a dependence of r^-1.64^. Interestingly, we found that the impedance of small electrodes with radius of 3 µm and 5 µm deviates from the linear fit of larger electrodes with radius greater than 10 µm. In this region, the impedance scales with a weaker dependence of the electrode radius (r^-0.83^). We confirmed that this deviation was not a fabrication error, by collecting SEM images of the smallest electrodes (figure 1d) and found that the electrode size is not significantly larger than designed.

**Figure 2.**
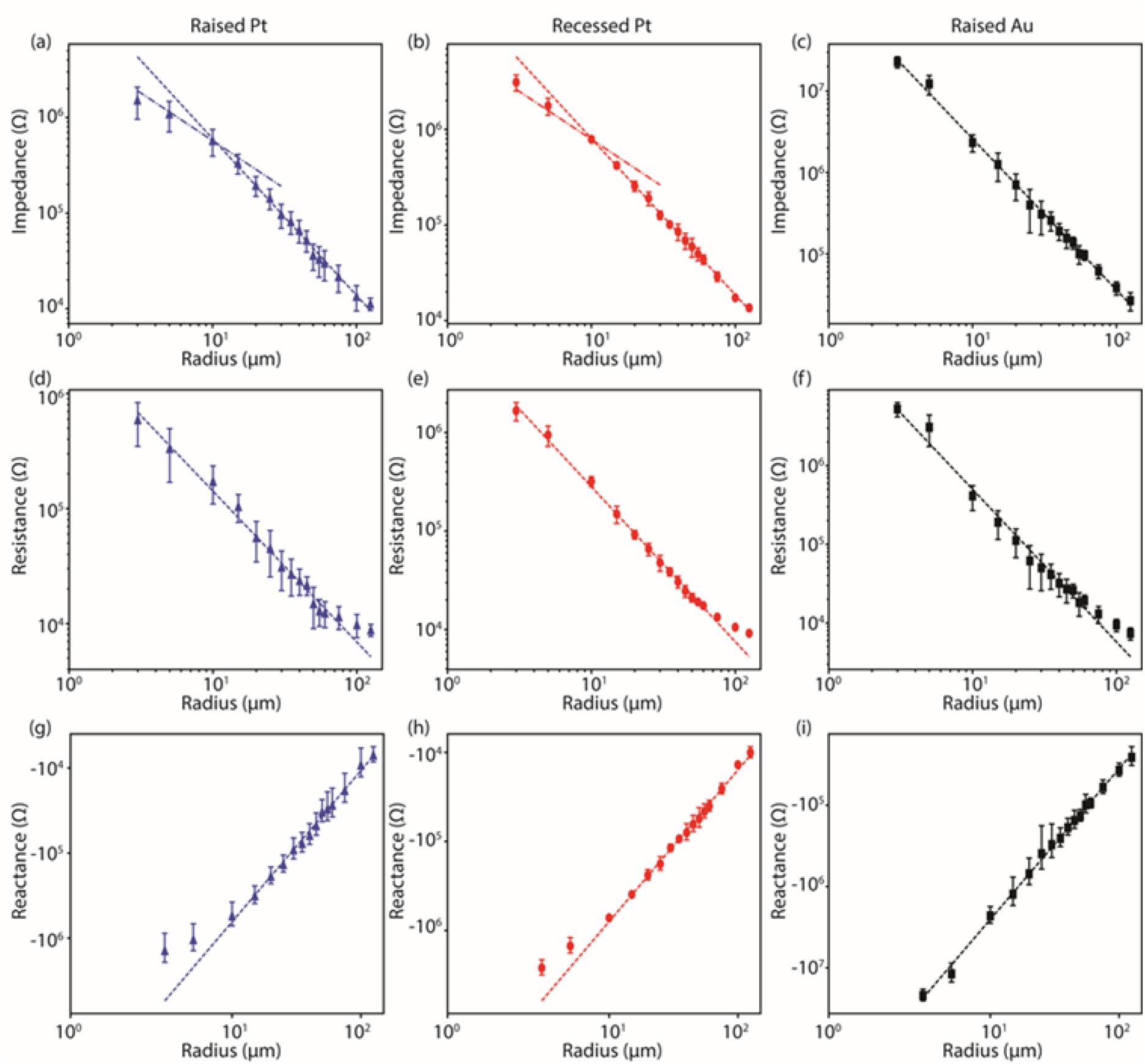
Impedance, resistance, and reactance at 1 kHz in log scale of raised Pt, recessed Pt and raised Au electrodes. (a)-(b) Corrected impedance of raised Pt (a) and recessed Pt (b) electrodes, slope of linear fits for all but two smallest electrodes are shown in dash line, a guideline with a slope of -1 is also shown (dot-dash line). (c) Corrected impedance of raised Au electrodes, slope of linear fit for all electrodes is shown in dash line. (d)-(f) Corrected resistance of raised Pt (d), recessed Pt (e), and raised Au (f) electrodes, slope of linear fits for all but three largest electrodes are shown in dash line. (g)-(h) Corrected reactance of raised Pt (g) and recessed Pt (h) electrodes, slope of linear fit for all but two smallest electrodes is shown in dash line. (i) Corrected reactance of raised Au electrodes, slope of linear fit for all electrodes is -1.86 (dash line). The slope values are shown in the Table 1.

We hypothesized that this deviation could be the result of the raised edges of the Pt electrode contributing to a larger electrode area, but recessed electrodes (figure 1b) also showed a similar low impedance. When we fabricated and tested electrodes with no exposed edges, we also found similar EIS (SI figure 1b,1e) and 1 kHz impedance across all the sizes (figure 2b). Linear fits to these data showed nearly identical scaling for large electrodes (>= 10 µm) as we had seen in the raised Pt electrodes (r^-1.64^). Smaller electrodes also showed a reduced dependence on radius (r^-1.17^), but not as significant as we observed for small, raised Pt electrodes.

**Table 1.** Slope of linear fit of impedance at 1 kHz for raised Pt, recessed Pt, and raised Au electrodes.

|  | Raised Pt | Recessed Pt | Raised Au |
| --- | --- | --- | --- |
| Impedance @ 1 kHz | -1.64 | -1.64 | -1.86 |
| Resistance @ 1 kHz | -1.31 | -1.57 | -1.94 |
| Reactance @ 1 kHz | -1.78 | -1.70 | -1.86 |

By examining the resistance (real part of the impedance) and reactance (imaginary part of the impedance) from our EIS data we determined that the deviation at small electrode sizes originates mainly from the reactance contribution (figure 2g, 2h). This can be seen by the fact that other than the two largest electrodes with radius of 100 and 125 µm, where series resistance dominates the total impedance^16^, the resistance fell on a linear fit for all electrode sizes, even for electrodes smaller than 10 µm (r^-1.31^ for raised Pt electrodes and r^-1.57^ for recessed Pt electrodes). On the other hand, the reactance shows a similar deviation in small electrodes as observed for total impedance: for larger electrodes, the reactance is dependent on r^-1.78^ for raised Pt electrodes and r^-1.70^ for recessed Pt electrodes.

Based on these data we hypothesized that this deviation is caused by a pseudocapacitance in combination with diffusion-controlled processes on the Pt electrode. Figure 3a shows a typical cyclic voltammogram of a microfabricated Pt electrode in PBS. At negative potentials around - 0.5 V vs. Ag/AgCl we observe hydrogen adsorption and desorption peaks. Beyond -0.6 V the onset of H_2_ evolution becomes evident. Around 0 V vs. Ag/AgCl we see a strong contribution of both, platinum oxide reduction and oxygen reduction, which are likely responsible for the observed pseudocapacitance. We tested this hypothesis by comparing impedance measurements to Au electrodes that do not exhibit such a pronounced pseudocapacitance around 0 V_ref_ and of which the impedance is mainly dominated by the double layer capacitance. For these experiments, we fabricated raised gold electrodes of the same design as the raised Pt electrodes (figure 1c). From the EIS measurements, we noticed that the impedance of small gold electrodes is in the range of 10 MΩ, which is comparable to the shunt impedance (also referred to as parasitic capacitance) of the system. The shunt impedance is caused by the capacitance and resistance between the traces of the multi-channel microelectrodes. For our probes, we measured a shunt impedance of 30.73 MΩ at 1 kHz (SI figure 2). This shunt impedance is in parallel with the electrodes, so we corrected the measured EIS data with this value using Ohm’s Law (see Supplementary information). The corrected impedance of raised gold electrodes at 1 kHz is shown in figure 2c, where the impedance has a dependence of r^-1.86^ for all the sizes without any deviation.

**Figure 3.**
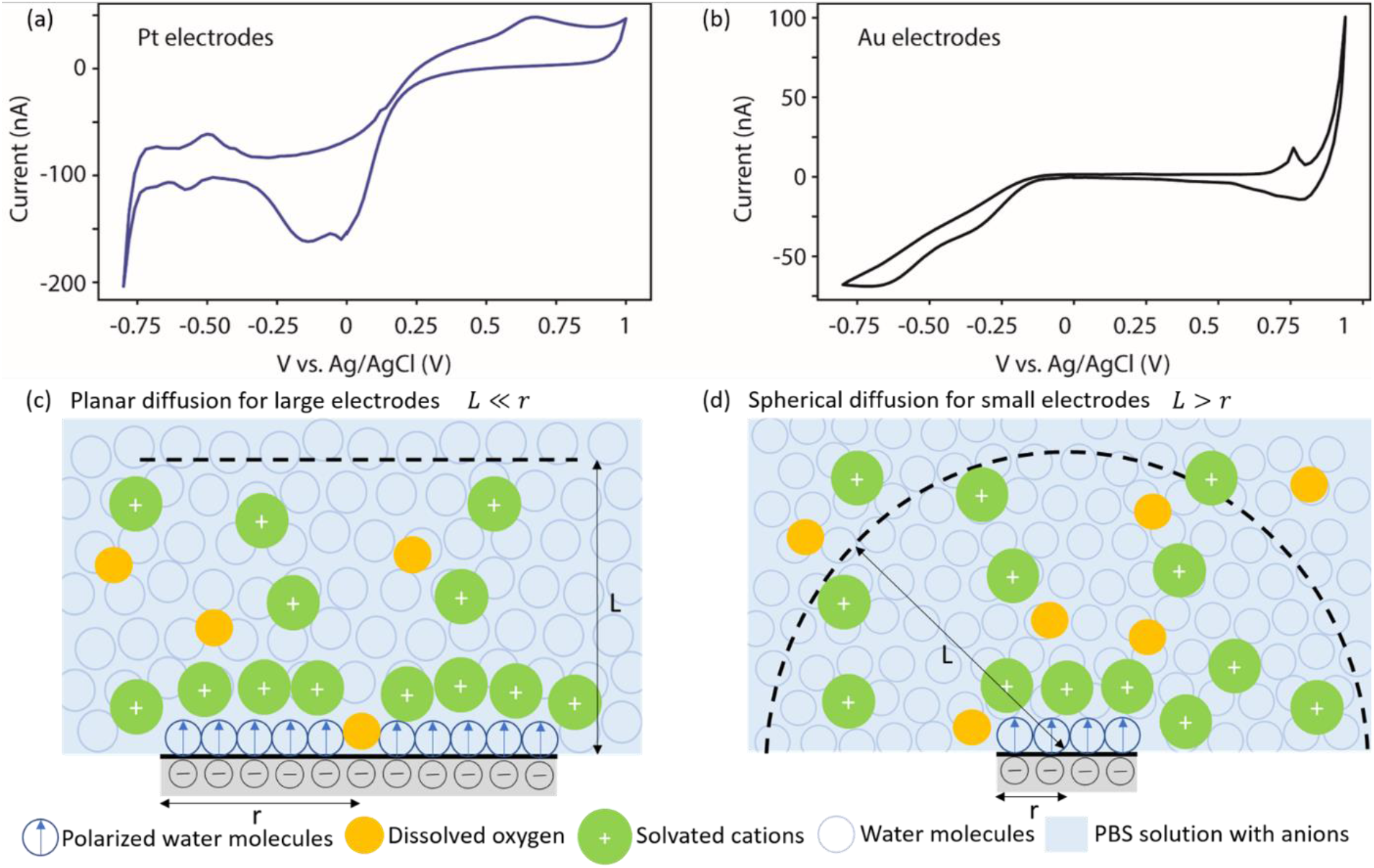
Cyclic voltammetry and diffusion model of Pt and Au electrodes. (a) Cyclic voltammetry of a Pt electrode in PBS, at 0 V vs. Ag/AgCl. (b) Cyclic voltammetry of a Au electrode in PBS, at 0 V vs. Ag/AgCl. (c) Schematic of linear and spherical diffusion profiles. For large electrodes, the boundary of the diffusion layer (L) is parallel to the electrode surface and the diffusion current has an area dependence of r^-2^. (d) For small electrodes (r < 10 µm), the boundary of the diffusion layer (L) is a half sphere, the diffusion current at steady-state is dependent on r^-1^.

A possible explanation for our observations can be found in early work on electrochemical characterization of microelectrodes^17^. As an electrode decreases in size there is a transition from a planar to a hemispherical diffusion profile for the reactants involved in the electrochemical reaction at the electrode surface. In our case, when the electrode (either platinum or gold) is large, the faradic current approaches a limit as described by the Cottrell equation^18^ with an area dependence (I ∼ r^2^) (figure 3c). As the electrode size becomes smaller than the thickness of the diffusion layer, the faradic current at steady state exhibits a radial dependence (I ∼ r) as described by the Saito equation^19^. As a result, the impedance changes scaling in the range from r^-2^, which is on the same order as a purely capacitive component at the interface, to r^-1^. This transition closely matches our observed scaling for the impedance of Pt electrodes that transitions from r^-1.6^ to r^-0.8^ for raised electrodes and r^-1.6^ to r^-1.2^ for recessed electrodes. The deviation from r^-2^ to r^-1.6^ is probably due to the existence of constant phase elements, which are influenced by surface roughness^20^ and non-uniform distribution of the surface R-Cs^21,22^. The spherical diffusion hypothesis is further supported by the difference between raised and recessed Pt electrodes in the anomalous impedance regime. Raised Pt electrodes show a larger effect (impedance ∼ r^-0.8^) than recessed Pt electrodes (impedance ∼ r^- 1.2^), which is expected due to the high SU8 passivation layer of 4 µm.

To confirm that the anomalous low impedance is related to diffusion we plot the impedance as a function of electrode size at frequencies of 1 Hz and 10 kHz. As expected, we find that at 10 kHz the impedance of Pt electrodes looks similar to Au electrodes and shows little deviation from the log-linear fit (r^-1.6^) (figure 4c, 4d). At 1 Hz, however, there is increased anomalous impedance scaling where electrodes with a radius of 10 µm begin to deviate from the log-linear fit for Pt. However, we see no deviation in impedance scaling for Au electrodes (figure 4a, 4b). These data are consistent with the hypothesis that a diffusion-limited process is the source for the anomalous impedance scaling of the Pt microelectrodes. At high frequencies we expect that the diffusion layer is small and will not dominate the impedance. For large electrodes, the impedance at high frequencies is dominated by the solution resistance with a r^-1^ dependence, as shown in figure 4c.

**Figure 4.**
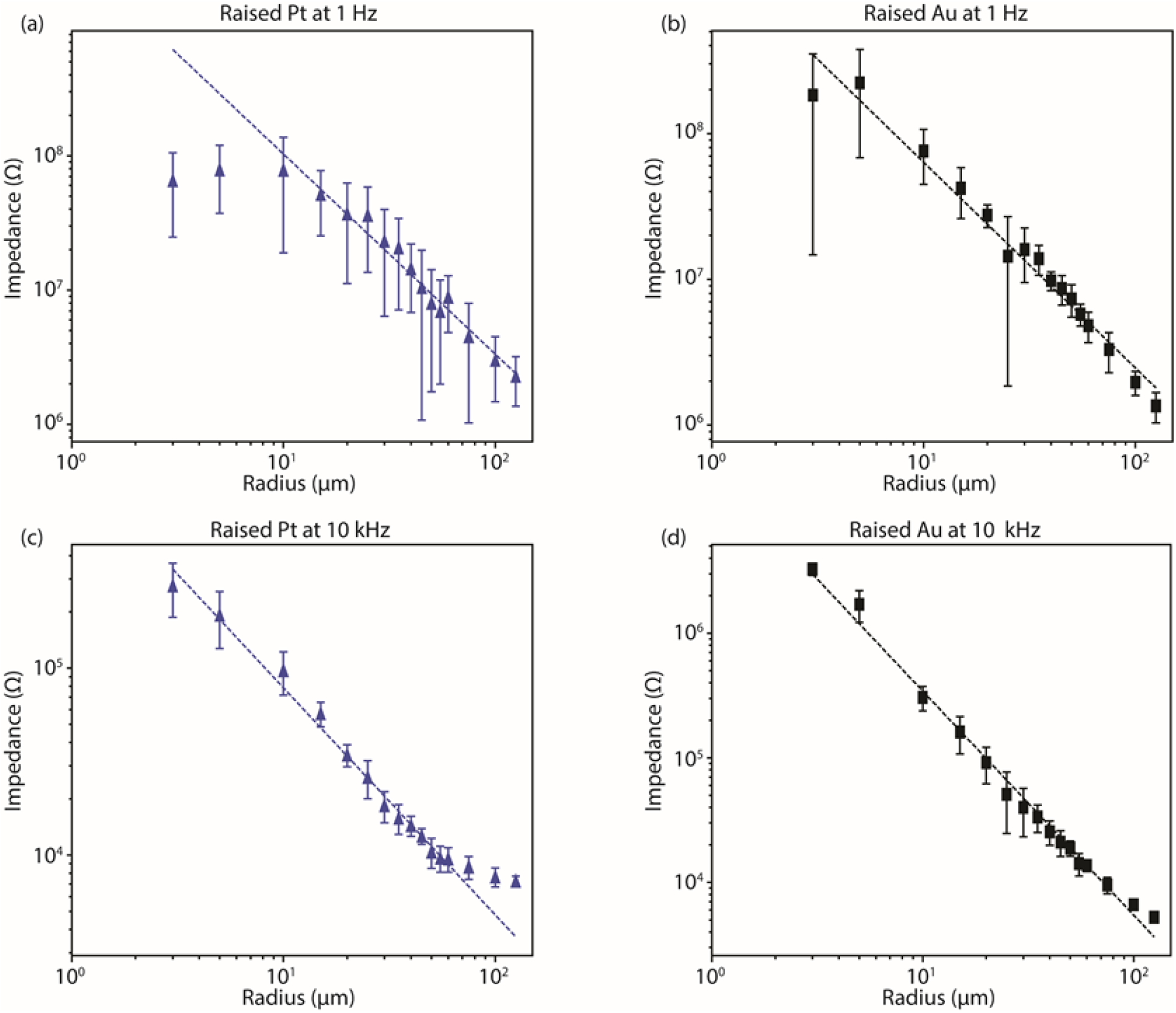
Impedance of raised Pt and raised Au electrodes at 1 Hz and 10 kHz. (a) Impedance of raised Pt electrodes at 1 Hz showing that for large electrodes with radius greater than 10 µm, the impedance scales with the dashed line (r^-1.49^), while for small electrodes, the impedance is dominated by reactance and is almost not related to the radius of electrodes. (b) Impedance of raised Au electrodes at 1 Hz scale as r^-1.41^ (dashed line) for electrodes of all sizes. (c) Impedance of raised Pt electrodes at 10 kHz scales as r^-1.21^ (dashed line) for electrodes of all but the three largest sizes. (d) Impedance of raised Au electrodes at 10 kHz scales as r^-1.80^ (dashed line) for electrodes of all sizes.

## Discussion

From our measurements we have found that for large Pt electrodes (r >= 10 µm) the impedance scales roughly with the electrode area (r^-2^) as previously reported^16,23^; however, as the electrodes fall below 10 µm radius their 1 kHz impedance deviates from this relationship, transitioning from approximately a r^-2^ scaling to an r^-1^ scaling. This change in scaling matches a transition from a planar to spherical diffusion layer which induces a current dependence transition from areal to radial. While diffusion-limited phenomena on disk microelectrodes were studied intensively in the electrochemistry field^17^, the development of neural microelectrode array devices has proceeded independently. Indeed, only when recording electrodes approach the micrometer regime these effects appear under physiological recording conditions.

By comparing Pt and Au electrodes, we confirmed that the transition in electrode scaling rates is caused by a diffusion layer that is most likely related to the oxide/oxygen reduction on the Pt surface. The transition from a planar to a spherical diffusion profile occurs when the electrode radius becomes small compared to the thickness of the diffusion layer. Because the thickness of this layer becomes larger at low frequencies, we see a more significant effect in our measurement of the 1 Hz impedance. As a result, as we measure how impedance scales based on electrode size and frequency we find a transition of radius dependence from r^-2^ to r^-1^ for Pt electrodes.

This finding of a change in impedance scaling for small Pt electrodes due to reactant diffusion is especially important for neural recording in the brain, where diffusion can be even more limited due to the neuropil. In this case, diffusion-limited processes may become significant even for larger electrodes than what we find in a PBS solution. Here we found that for ultra-small Pt microelectrodes, at a critical size of radius less than 10 µm, 1 kHz impedance of electrodes made by materials with pseudo-capacitance is weakly related to electrode size.

These results can also explain why fractal electrodes or waffle shaped electrodes showed reduced impedance compared to planar electrodes of a similar area due to an “edge effect”^12^.

An open question is how this divergence in impedance measurements affect neural electrode performance in vivo. In stimulation experiments, it is essential to have low impedance electrodes to prevent the requirement of high voltage application for driving the necessary current. However, for recording, it has been argued that the common target of 100 kΩ impedance at 1 kHz is not necessarily an important metric for high quality neural recordings given modern high-impedance neural amplifiers^13^. Indeed, if the electrode impedance is below the input impedance of the amplifier one would not expect signal attenuation provided there was sufficiently small stray capacitance in the electrode insulation^11,13^. A reduced electrode impedance may help the most for extremely small electrodes where one would expect thermal noise to otherwise overwhelm the signal amplitude. It remains to be seen if ultra-small Pt electrodes can indeed perform well for chronic neural recordings; however, promising results have been obtained in vitro for nano-scale Pt electrodes giving some prospect for similar performance in future in vivo probes^24,25^.

## Methods

### Fabrication

All the electrodes are fabricated using a similar method as previously reported^12^. We first sputtered a release layer of 70 nm Al on a Si wafer (University Wafer). Followed by photolithography to pattern the bottom SU8: spin coated SU8 2005 (MicroChem) at 5000 rpm for 30 s, resulting in a 4 µm thick layer, then prebaked at 65 °C and 95 °C for 1 min and 4 min, respectively. After exposure, the wafer was post-baked again at 65 °C and 95 °C for 1 min and 4 min, respectively, then developed in SU8 developer for 2 min. Next, LOR 3A was spin-coated at 2000 rpm for 1 min, baked at 100 °C and 150 °C for 2 min and 10 min as lift-off sacrificial layer. Then S1818 was spin-coated at 4000 rpm for 1 min, baked at 115 °C for 1 min. After exposure, the wafer was developed in MF321 for 2 min. Prior to sputtering metal traces, the surface was treated with oxygen plasma at 100 W for 1 min to enhance adhesion. A 100 nm thick Pt layer was sputtered and lifted off with acetone. For gold electrodes, 8 µm-thick Ti was sputtered as an adhesion layer. Next, top insulation SU8 was patterned using the same steps as bottom SU8. For recessed electrodes, a dry etch was applied using Reactive Ion Etcher (Oxford 180) to open up the top SU8 vias for small electrodes. Before releasing, the microelectrodes were hard-baked at 180 °C for 1 hour. The microelectrodes were released using MF 321 for 8 hours.

### Electrical connection and electrical measurements

After releasing, the microelectrodes were collected and soaked in DI water for 24 hours. For EIS measurements, the electrodes were connected to break-out boards (OSH Park) using silver print (Silver Print II, GC Electronics). The EIS was performed with a three-electrodes setup using a Gamry 600 + electrochemical station. The reference electrode was a silver-silver chloride (Gamry Instrument) standard electrode filled with saturated potassium chloride, and the counter electrode was a bare Pt wire (0.010’’ in diameter, uGems). The electrodes emerged in 1X phosphate buffered saline (PBS) (Fisher). The EIS was measured from 100 kHz to 1 Hz, with 30 points per decade. The input voltage was 10 mV in root-mean-square.

For cyclic voltammetry measurements, we used the same three electrodes set up as during EIS measurements, a Pt electrode and a Au electrode with a radius of 120 µm were picked as working electrodes. For both scans, voltage was scanned from -0.8 V ∼ 1 V vs. reference electrode, with a scan rate of 200 mV/s.

## Acknowledgments

We thank Rice Shared Equipment Authority for fabrication support. This work was funded by NIH BRAIN R21 to JTR.

