## Supplementary information for "Impedance scaling for gold and platinum microelectrodes" (from another version of this preprint: version 1, 2021-03-10)

**Impedance correction:**

All impedance, resistance and reactance in figure 3 and figure 5 were corrected using Ohm's Law:

$$\frac{1}{Ztotal} = \frac{1}{Zshunt} + \frac{1}{Zelectrode}$$

Where Z = Zreal + i*Zimag, all computation was complex. For impedance at 1 kHz, 1 Hz and 10 kHz, the impedance was corrected using shunt impedance at 1 kHz, 1 Hz and 10 kHz, respectively.

**Simulated reactance:**

For large electrodes:

$C = \frac{\varepsilon_0 \varepsilon_r A}{\mathrm{d}}$, where $\varepsilon_r$ = 80, with $\varepsilon_0$ = $8.85 \times 10^{-12}$, d = 0.5 nm.

For smaller electrodes:

$C_{\ half\ sphere} = \frac{\mathrm{k} \times 4\pi \varepsilon_0 \varepsilon_r}{2(r^{-1} - L^{-1})}$, where $\varepsilon_r$ = 80, with $\varepsilon_0$ = $8.85 \times 10^{-12}$, L = 20 µm, k is a constant of 5000 to make the simulated reactance using both equations have the same value at r = 10 µm.

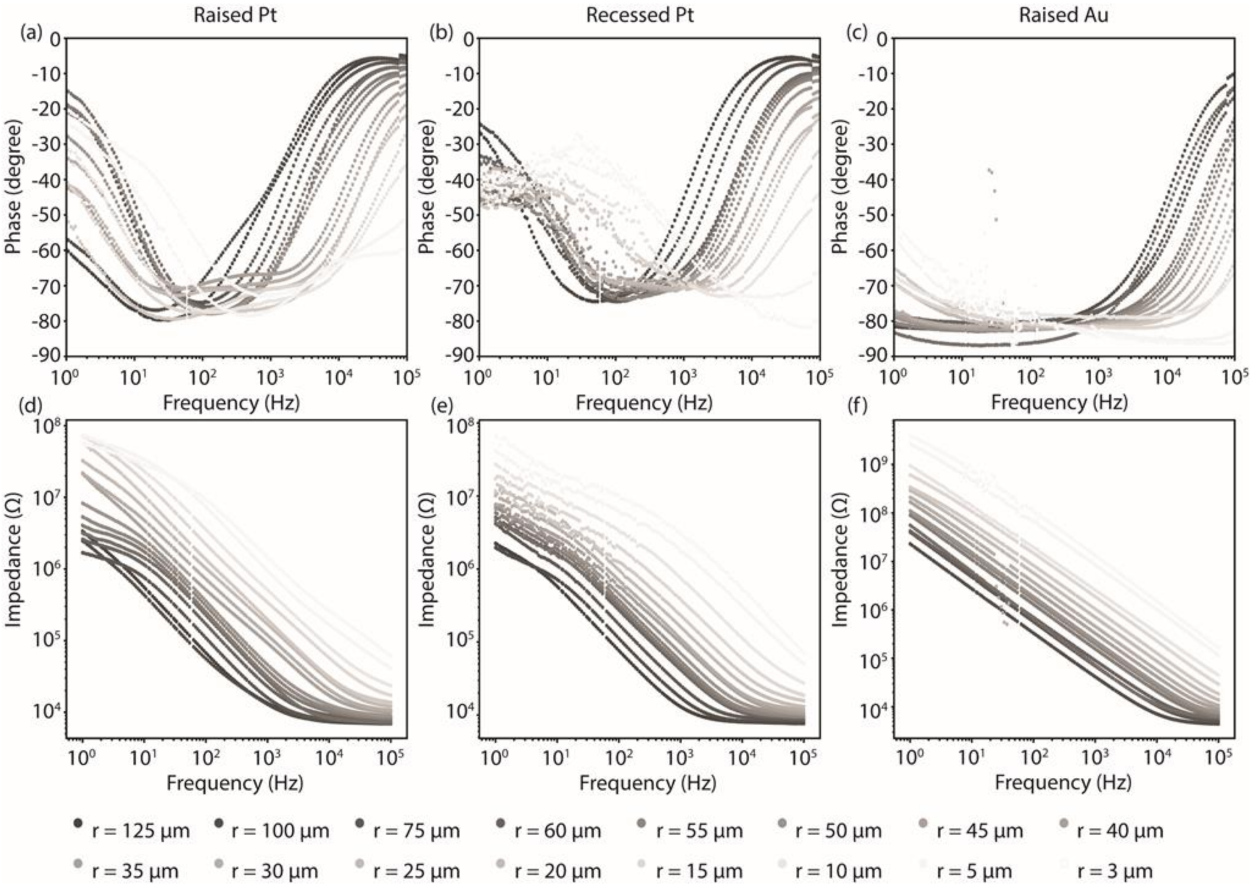


**SI Figure 1. Bode plots of raised Pt, recessed Pt and raised Au electrodes.** (a)-(c) Phase of raised Pt, recessed Pt and raised Au electrodes, respectively. (d)-(f) EIS of raised Pt, recessed Pt and raised Au electrodes, respectively. The measurement was unstable at low frequencies, some data points jumped around.

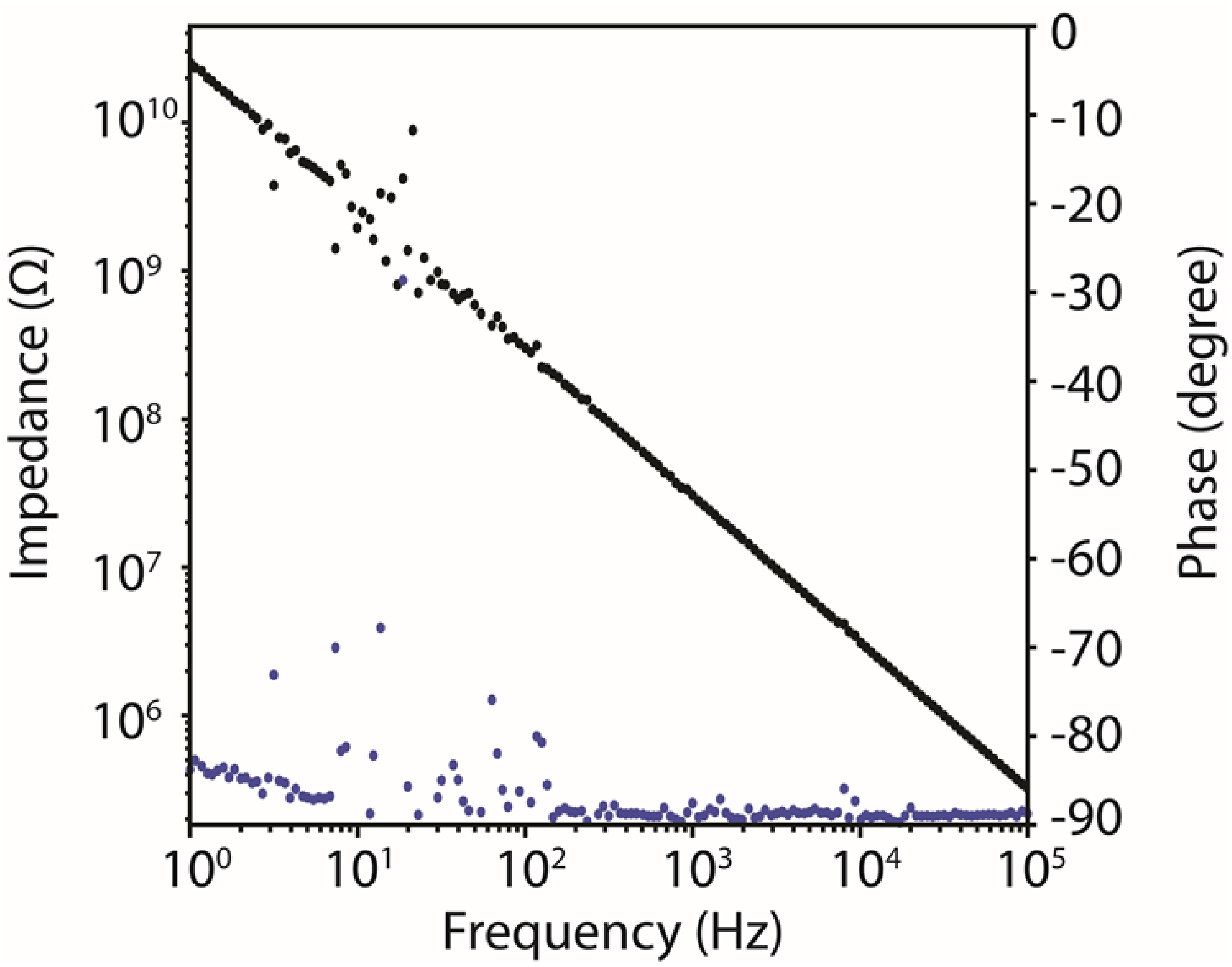


**SI Figure 2. Bode plot of shunt impedance.** The impedance is shown in black and the phase is shown in blue. The shunt impedance is mostly shunt capacitance at high frequencies. The measurement between 10-100 Hz was unstable.
